# Early diet programs feeding circuits and food solicitation behavior

**DOI:** 10.64898/2026.03.28.714959

**Authors:** Marie-Therese Fischer, Camilo Rodriguez Lopez, Billie C. Goolsby, Max Madrzyk, Ling-Yi Zhang, Lauren A. O’Connell

**Affiliations:** Department of Biology, Stanford University, Stanford, CA 94305, USA; Department of Microbial Ecology, University of Vienna, 1030 Vienna, Austria; Wu Tsai Institute for Neuroscience, Stanford University, Stanford, CA 94305, USA

**Keywords:** urocortin, CART, begging, risk avoidance, aggression, artificial diet, natural diet, phosphoTRAP

## Abstract

From birth, offspring must balance internal energy state with the costs of signaling need to caregivers to secure nutrition for growth and healthy development. How diet during this period shapes the developing brain and behavior is a central question in biology, with growing relevance for understanding the developmental origins of metabolic disorders such as obesity. Yet the effects of early nutrition on growth, neural development, and behavior are rarely examined together within a single system. Here, we use poison frog tadpoles as a vertebrate model in which offspring develop outside a womb, can be reared independently under precisely controlled dietary conditions, and display diverse behaviors at a young age. By independently manipulating dietary quantity and quality, we link early nutrition to growth, brain development, and social behavior during ontogeny. Diet quantity increased body size, whereas diet quality accelerated development. Diet quality, but not quantity, shaped behavior: tadpoles fed a natural diet showed increased affiliation during food solicitation and reduced feeding, while aggression and risk avoidance remained unchanged. Diet also affected body and brain development differently, with artificial diets producing larger tadpoles but reduced volume in some brain regions, including areas associated with begging behavior. Anorexigenic urocortin-1 neurons were active during begging behavior and, together with another anorexigenic midbrain population, decreased in abundance under an artificial diet. Functionally, urocortin-1 reduced feeding without directly increasing begging, suggesting it gates socio-positive behavior by suppressing a competing drive rather than initiating it. Together, this work shows that dietary quantity and quality act on distinct developmental processes, with diet quality programming the brain circuits that permit begging behavior while growth is determined by food quantity.

## Introduction

Adequate nutrition during early development is essential for healthy growth and lifelong wellbeing (1–8). As infants must balance internal energy demands with the energetic costs of communicating hunger to caregivers, nutritional state and food-solicitation behavior are tightly coupled from the beginning of life (9–13). While affiliative signaling of need is the primary behavioral expression of hunger in many altricial infants, nutrition influences a broader spectrum of behaviors across vertebrates, including aggression, foraging decisions, and responses to risk. For example, competition over limited food resources can elicit aggression among siblings across taxa (14–18), and nutritional state can shift how individuals balance food acquisition against predation risk, with food deprivation predisposing individuals to high-risk behaviors (19–21). Together, these findings suggest that early-life diet shapes not only growth but also neural circuits that link nutritional state to behavior.

The neural mechanisms that integrate energy state with behavioral decision-making remain poorly understood. Work in rodents has identified neuromodulatory systems that regulate appetite and feeding in adults, including neuropeptide Y (NPY), pro-opiomelanocortin (POMC), cocaine-and amphetamine-regulated transcript (CART), agouti-related protein (AgRP), orexin, galanin, urocortin and basonuclin neurons (22, 23). These circuits are organized prenatally (24–26) and are used to assess energy balance and regulate feeding. They are well positioned to integrate internal energetic state with external cues to guide behavioral prioritization when animals face competing demands (9, 27). However, their function can differ markedly between early life and adulthood. For example, ablation of AgRP neurons causes starvation in adult mice but has no effect in neonates (28), where these neurons respond to social isolation rather than caloric deprivation (29). Such findings suggest developmental differences in how feeding-related circuits regulate behavior, highlighting the need to evaluate these networks not only in adults but also during early life. Nevertheless, how prenatal diet shapes the wiring of these circuits, and how such changes translate into measurable social behaviors, has rarely been quantified in young animals. Addressing this question in mammals is challenging due to in utero brain development, prolonged maternal dependency, and limited behavioral repertoires in neonates that are largely restricted to caregiver-directed affiliation.

To overcome these obstacles, we address this question in poison frog tadpoles, where offspring can be reared independent from parents and diet can be precisely controlled (30, 31). Amphibians in general are a promising model system for studying brain organizational principles as the same brain regions and neuropeptides that regulate feeding and social behavior in mammals are conserved and readily apparent in amphibians, enabling generalizable insights from a simple nervous system (32, 33). In the mimic poison frog (*Ranitomeya imitator)*, parents form monogamous pairs and jointly rear offspring (15, 34). After hatching, fathers transport the cannibalistic tadpoles individually to small bodies of water that form in leaf axils and are scarce in nutritional resources (35). Every few days, the mother returns to feed her tadpoles by depositing trophic unfertilized eggs into the nursery (36), a physiological production of food analogous to mammalian lactation that constitutes their primary food source. Limited resources prime tadpole social interactions and can manifest in two distinct behaviors: an affiliative interaction by displaying a conspicuous begging behavior to a caregiver or an agonistic interaction by displaying aggression towards conspecific larvae (36, 37). In tadpoles, begging behavior is an honest signal for hunger (38), and is not attributable to social distress or thermoregulatory need like in rodent pups (39, 40). Beyond these social interactions, nutritional state also drives independent foraging on plant material (37) and may influence decisions to prioritize feeding over remaining immobile under predation risk (41). This system provides a powerful framework to test the hypothesis that early-life nutrition programs the development of neural circuits that sense internal energy state and integrate this information into behavioral decision-making. Specifically, we predicted that dietary quality and quantity would influence the development of appetite-regulating neuronal populations and, in turn, modulate behaviors such as food solicitation, aggression, feeding, and risk avoidance.

## Results

### Diet shapes developmental rate and body size

We assigned *R. imitator* tadpoles to five dietary treatments designed to independently manipulate food quality and quantity (Fig. 1A, D). Four groups received an artificial diet based on a commercially available tadpole food and half these groups were supplemented with brine-shrimp flakes to generate higher-quality food. Two groups received a low-quantity regimen with four times less food, thereby generating a two-by-two artificial diet experiment that varied in quantity and quality. As a natural reference, one group of tadpoles received two frog eggs per week, approximating parental provisioning in the wild (37).

**Figure 1.**
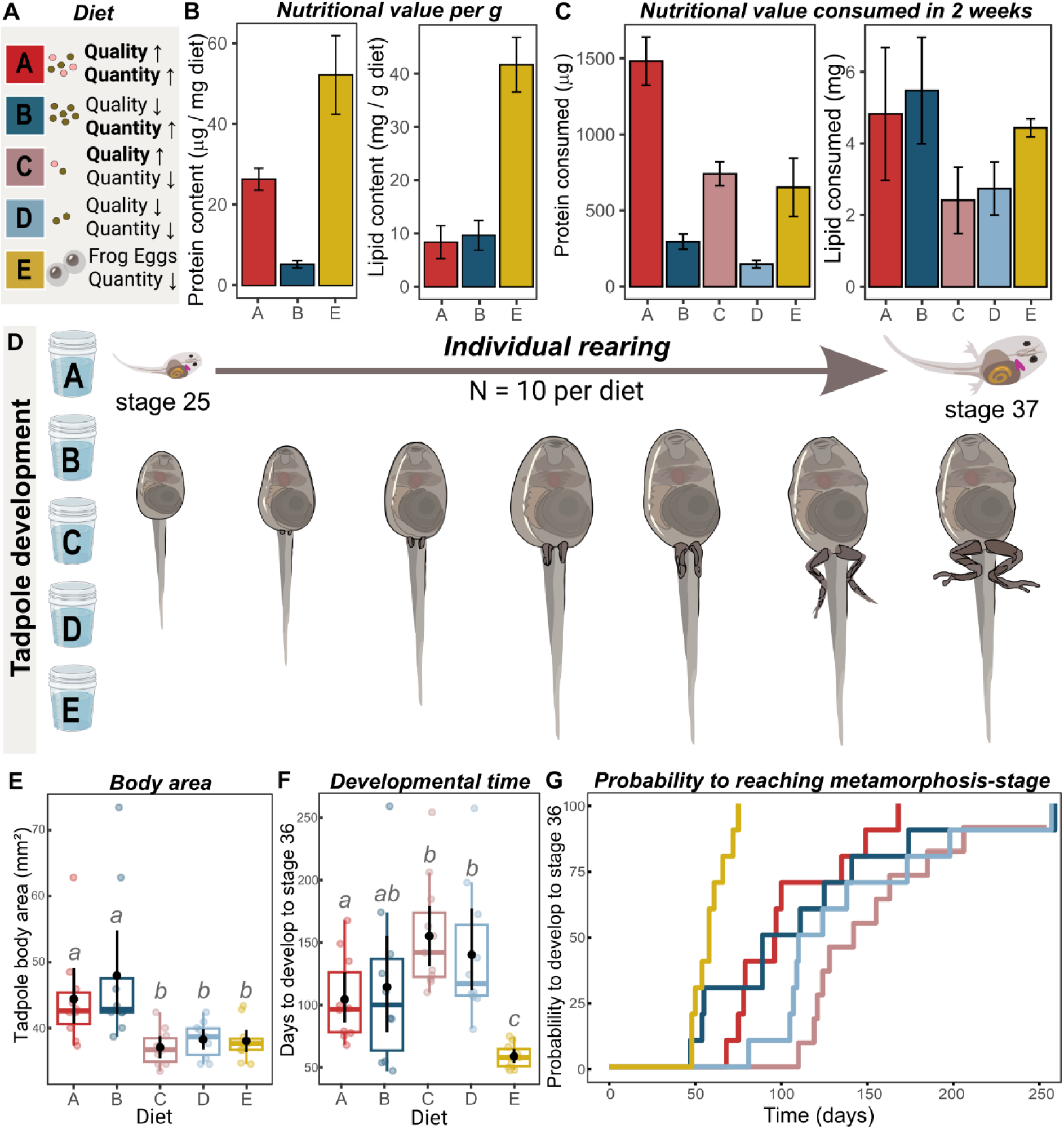
Developmental consequences of different nutritional regimes. **(A)** Overview of five dietary treatments manipulating food quality and quantity: enriched artificial diets (A, C), base artificial diets (B, D), and a natural egg diet (E). Diets A and B were provided in high quantity, diets C and D in low quantity, and diet E consisted of two frog eggs per week. **(B)** Protein (left) and lipid (right) content per gram of diet, quantified by Bradford assay and gravimetric lipid extraction (N = 4 per diet). **(C)** Total protein (left) and lipid (right) content over two weeks, accounting for differences in feeding quantity. **(D)** 10 tadpoles per diet were reared in individual cups and measured and staged weekly based on limb/toe development until they reached stage 36-37 (42) **(E)** Tadpole body area at matched developmental stages; different letters indicate significant post-hoc differences (Kruskal–Wallis test, adjusted p < 0.05). **(F)** Developmental time to stage 36 (N = 10 per diet); black dots depict mean values with 95% confidence intervals, boxplots show median and interquartile range, with letters denoting significant differences. **(G)** Kaplan–Meier curves showing the probability of reaching Gosner stage 36 over time.

Quantification of lipid and protein content revealed marked differences in macronutrient composition among diets (Fig. 1B). On a per-gram basis, the egg-diet contained the highest levels of both lipids and proteins, followed by the enriched artificial diet. When accounting for differences in feeding quantity over two weeks, enriched high quantity diet provided the highest total protein amount and base high quantity diet the highest total lipids, while the egg-diet closely resembled the high quantity/rich diet in total lipids and the low quantity/rich diet in total protein (Fig. 1C).

We expected that if total food quantity primarily drove developmental and behavioral outcomes, differences would align with quantity treatments. By contrast, if food quality were more influential, we predicted that differences would either arise between enriched and base artificial diets if they were mediated by protein content, or between artificial and natural feeding regimes if they were mediated by factors other than macronutrient content.

Tadpole body area differed significantly among diets (Kruskal–Wallis χ²₄ = 22.41, p < 0.001) (Fig. 1E). Tadpoles reared on high-quantity diets (A, B) were significantly larger than those on low-quantity diets (C– E; adjusted p < 0.05). Generalized linear modeling revealed a strong effect of diet quantity on body size (estimate = –0.190 ± 0.038, z = –4.95, p < 0.001), whereas dietary quality had no significant influence (p > 0.1; Fig. S1A). On average, tadpoles on high-quantity diets reached approximately 21 % larger body sizes at identical developmental stages than those on low-quantity diets, indicating that food quantity, rather than quality, determined body growth.

Developmental time differed significantly among diet treatments (likelihood ratio test = 25.89, df = 4, p < 0.001) (Fig. 1F, Fig. S1B). Development was slowest under low-quantity diets and fastest under the natural diet, with all other diets showing significantly reduced developmental rates (HR = 0.011–0.092, p < 0.05; Fig. 1G). When testing dietary predictors, low food quantity reduced the probability of metamorphosis (HR = 0.33, p = 0.019), whereas natural diet quality strongly accelerated development (HR = 65.51, p < 0.001).

### Diet quality shapes begging and feeding behavior but not aggression and risk avoidance

To assess how developmental diet affected behavior, tadpoles were tested in four behavioral paradigms. Each tadpole was presented with a female frog to assess socio-positive food-soliciting behavior (“begging” hereafter), food to measure feeding motivation, a smaller conspecific tadpole to quantify aggression, and food presented together with a predator to assess behavioral inhibition (Fig. 2A). This last behavior test is a novel risk-avoidance assay that captures suppression of movement and feeding when food was presented alongside a life predator, relative to food alone (movement: V = 330, p = 0.002; feeding: V = 223, p < 0.001) (Fig. S2).

**Figure 2.**
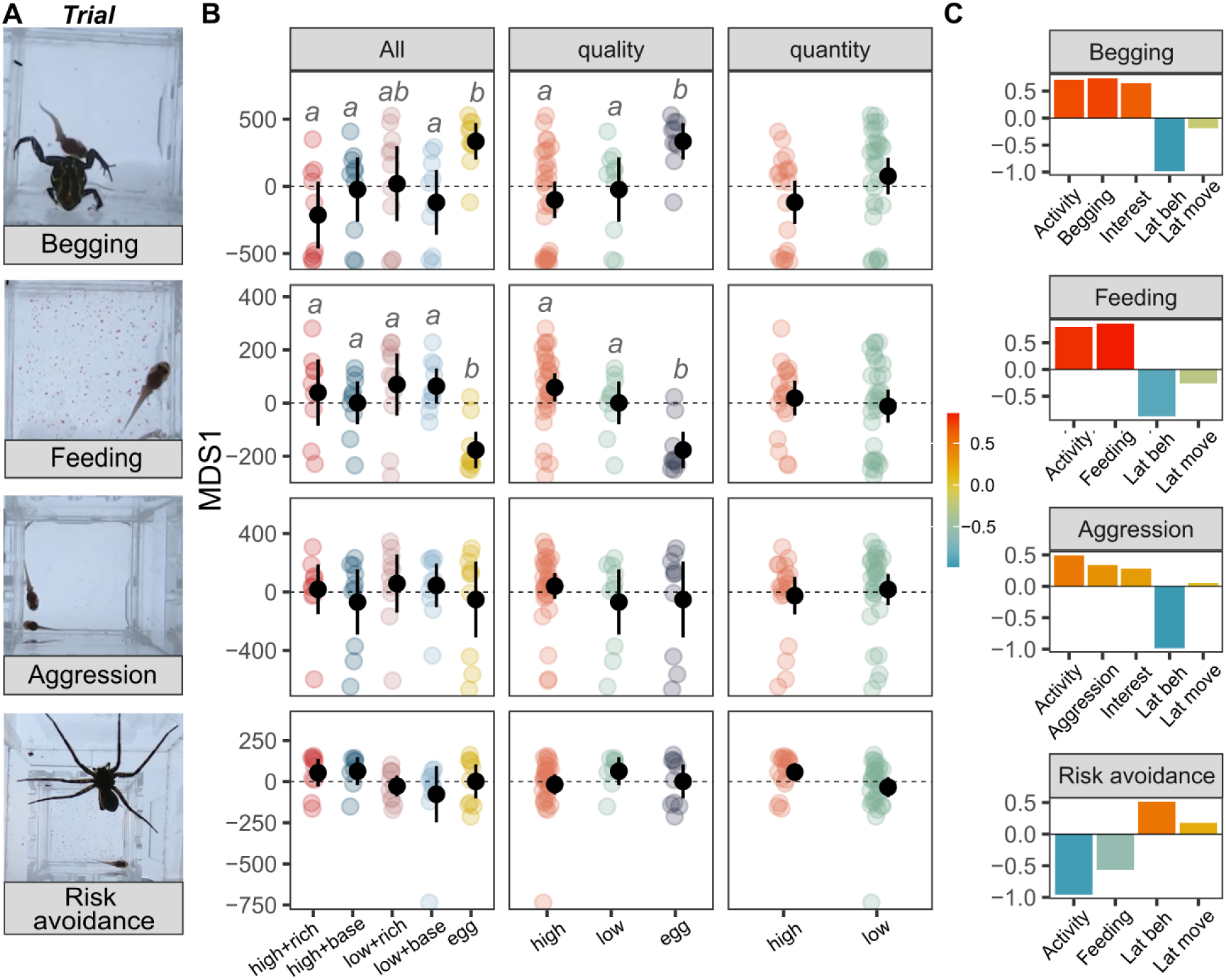
Developmental diet shapes socio-positive and feeding behavior but not aggression or risk avoidance. **(A)** Behavioral paradigms used to quantify begging (female frog), feeding motivation (food), aggression (smaller conspecific), and risk avoidance (food presented with a predator). **(B)** Metric multidimensional scaling (mMDS) of behavioral differences among rearing diets for each behavioral context. Points represent individuals; black dots indicate group means ± 95% confidence intervals. Panels show effects of individual diets (left), diet quality (middle), and diet quantity (right); different letters denote significant post-hoc differences following PERMANOVA (adjusted p < 0.05). **(C)** Variable loadings on the first mMDS axis (MDS1) for each trial, showing contributions of activity, begging/aggression, interest, and latency measures. Abbreviations: Lat beh = latency to behave; Lat move = latency to move.

Developmental diet significantly affected begging (PERMANOVA: F₄,₄₈ = 3.15, R² = 0.22, p = 0.02) and feeding behavior (F₄,₄₈ = 3.97, R² = 0.26, p = 0.001), but not aggression (F₄,₄₈ = 0.63, R² = 0.05, p = 0.74) or risk avoidance (F₄,₄₈ = 1.11, R² = 0.09, p = 0.33) (Fig. 2B). Specifically, egg-fed tadpoles showed significantly higher socio-positive food solicitation than tadpoles on diets high quantity and low quantity base diets (p.adj < 0.02), with a similar trend relative to low quantity enriched diet (p.adj = 0.08), and significantly reduced feeding activity compared to all other diets (p < 0.01) (Fig. 2B). Grouping treatments by diet quality and quantity revealed that quality, but not quantity, significantly influenced begging and feeding behavior (begging: F₂,₅₀ = 5.09, p = 0.005; feeding: F₂,₅₀ = 7.09, p = 0.002), whereas neither parental relatedness nor trial order explained behavioral variation (PERMANOVA, F ≤ 1.9, p > 0.1). Specifically, behavior differences were driven by contrasts between the natural and artificial diets, whereas behavior did not differ between base and enriched artificial diets. We assessed the consistency of begging behavior by re-testing each tadpole three days after the initial trial, revealing repeatable individual behavior (R = 0.37, p < 0.01; Fig. S3).

### Urocortin-1 neurons link diet and begging behavior

To investigate the neural basis of diet-dependent behavioral differences, we first characterized activity of nutrition-sensing neuronal populations during behaviors that were increased (socio-positive) or unaffected (aggression), relative to a non-social handling control (Fig. 3A). As egg-fed tadpoles showed reduced feeding but increased begging behavior, we hypothesized that diet-dependent behavioral modulation involves neuropeptidergic populations that regulate internal motivational states, possibly by suppressing appetite. We therefore predicted increased activity and potentially abundance of anorexigenic (appetite suppressing) neurons, or decreased activity or abundance of orexigenic (appetite stimulating) neurons.

**Figure 3.**
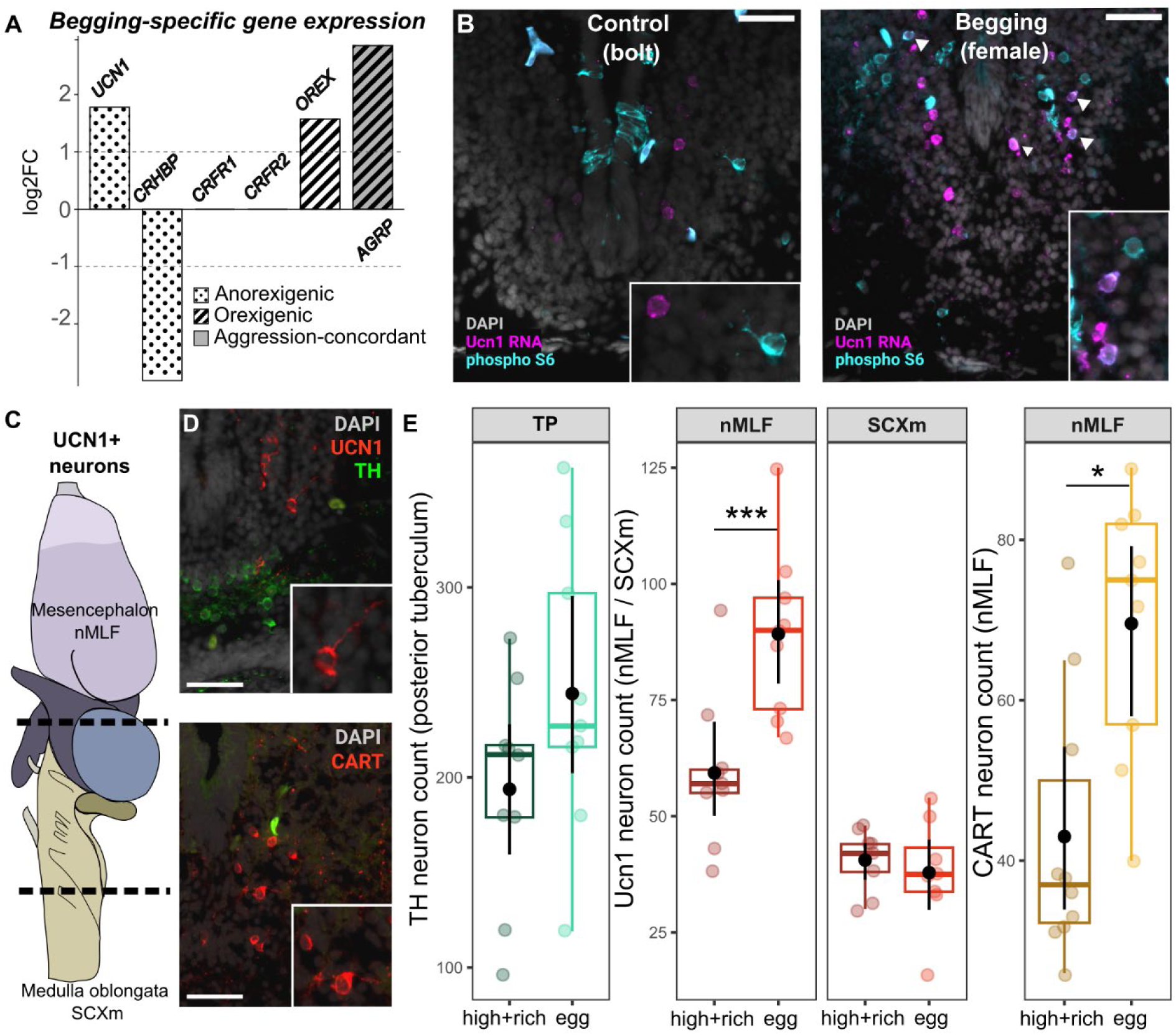
Ucn1-positive feeding-suppressing neurons are active during begging and more abundant in the brains of tadpoles fed a natural diet. **(A)** Begging-specific gene expression identified by phospho-TRAP (pTRAP; N = 5 per behavior), shown as log₂ fold change of phosphorylated S6–associated RNA relative to total input. Genes with absolute log₂ fold change > 1 and adjusted p < 0.1 were considered significantly altered and classified as anorexigenic or orexigenic; genes also regulated during aggression are shown in gray. **(B)** Double labeling of Ucn1 mRNA (isHCR) and phosphorylated ribosomal protein S6 (pS6) in handling control (bolt) and begging (female) conditions; arrowheads indicate UCN1⁺/pS6⁺ cells. **(C)** Lateral schematic of the tadpole brain showing two major Ucn1⁺ neuron locations along the anteroposterior axis (dashed lines) **(D)** Representative immunohistochemistry staining for Ucn1/TH, and CART on consecutive cryosections. **(E)** Neuron counts of TH⁺ cells (TP), Ucn1⁺ cells (nMLF, SCXm) and CART⁺ cells (nMLF) in tadpoles reared on enriched artificial diet or natural diet (N = 10 per group). Points represent individuals; black dots indicate group means ± 95% confidence intervals; boxplots show median and interquartile range, asterisk indicate significant differences.

To identify candidate populations, we used PhosphoTRAP (43) to identify signatures of translationally active neurons associated with begging but not aggressive behavior. Among the enriched gene set, urocortin-1 (Ucn1) showed the strongest selective enrichment during begging behavior (1.8 log₂ fold change), but not during aggression, identifying it as a key candidate in facilitating socio-positive behavior (Fig. 3A). In addition, transcription of corticotropin-releasing hormone-binding protein CRHBP, which binds urocortin-1 with high affinity, was reduced during socio-positive behavior (−3 log₂ fold change) while no transcriptional enrichment of the canonical receptors CRFR1 or CRFR2 was detected. This pattern is consistent with increased availability of urocortin-1 peptide without concomitant changes in receptor expression. We confirmed that Ucn1-expressing neurons were active during socio-positive food solicitation but not under control conditions (25% versus 1.4% of Ucn1^+^ cells active) (Fig. 3B). PhosphoTRAP analyses further revealed behavior-specific regulation of additional neuromodulatory receptor systems that were not examined further in the present study (Table S1, S2), suggesting broader circuit-level modulation beyond currently described dopaminergic pathways (44).

Building on the finding that anorexigenic Ucn1 neurons are active during begging we next examined whether developmental diet influenced the abundance of Ucn1-expressing neurons in tadpole brains. We focused on the two diet groups showing the strongest behavioral divergence: tadpoles fed a high-quality artificial diet and those fed an egg-diet. Neurons with high Ucn1 expression were restricted to two motor-associated nuclei: the mesencephalic nucleus of the medial longitudinal fascicle (nMLF) and the spinal cord nucleus motorius nervi vagi (SCXm) (Fig. 3C, D, Fig. S4A).

Ucn1 neuron abundance varied with diet and brain region (nutrition: χ²₁ = 4.85, p = 0.028; brain region: χ²₁ = 433.72, p < 0.001; nutrition × brain region: χ²₁ = 13.27, p < 0.001), while age had no effect (χ²₁ = 0.58, p = 0.45) (Fig. 3E). Nutritional effects on Ucn1 neuron abundance were region specific, with increased Ucn1 neuron numbers in the midbrain (nMLF: z = –3.70, p = 0.0002) but no diet effects in the hindbrain (SCXm: z = 0.11, p = 0.92).

To distinguish general diet-effects on anorexigenic systems from targeted modulation of begging circuits, we analyzed CART-expressing neurons in the nMLF and TH-expressing neurons in the posterior tuberculum (TP), a population required for begging behavior in *R. imitator* tadpoles (44) (Fig. 3E). CART neuron abundance differed significantly between diets (χ²₁ = 6.09, p = 0.014), with higher counts in egg-fed tadpoles but no effect of age (χ²₁ = 2.07, p = 0.15) (Fig. 3E). In contrast, TH neurons showed a modest increase across dietary conditions but were not significantly affected by diet (χ²₁ = 0.083, p = 0.77) or age (χ²₁ = 0.52, p = 0.47) (Fig. 3E). Together, these results show that nutritional quality robustly affects appetite-suppressing neuron populations, but not a neuropeptidergic population required for begging behavior in tadpoles.

### Dietary quality affects body and brain development differentially, with region-specific effects in brain areas associated with begging behavior

As begging behavior may scale with the absolute size of relevant brain regions rather than with selective expansion of specific neuronal populations, we tested whether dietary quality affected growth of brain regions associated with begging behavior (Fig. 4A, B). At stage 37, egg-fed tadpoles had significantly greater brain length along the body axis (t₁₆ = 3.01, p = 0.008) but reduced body size compared to tadpoles fed diet A (LR χ²₁ = 5.00, z = -2.24, p = 0.025) (Fig. 4A).

**Figure 4.**
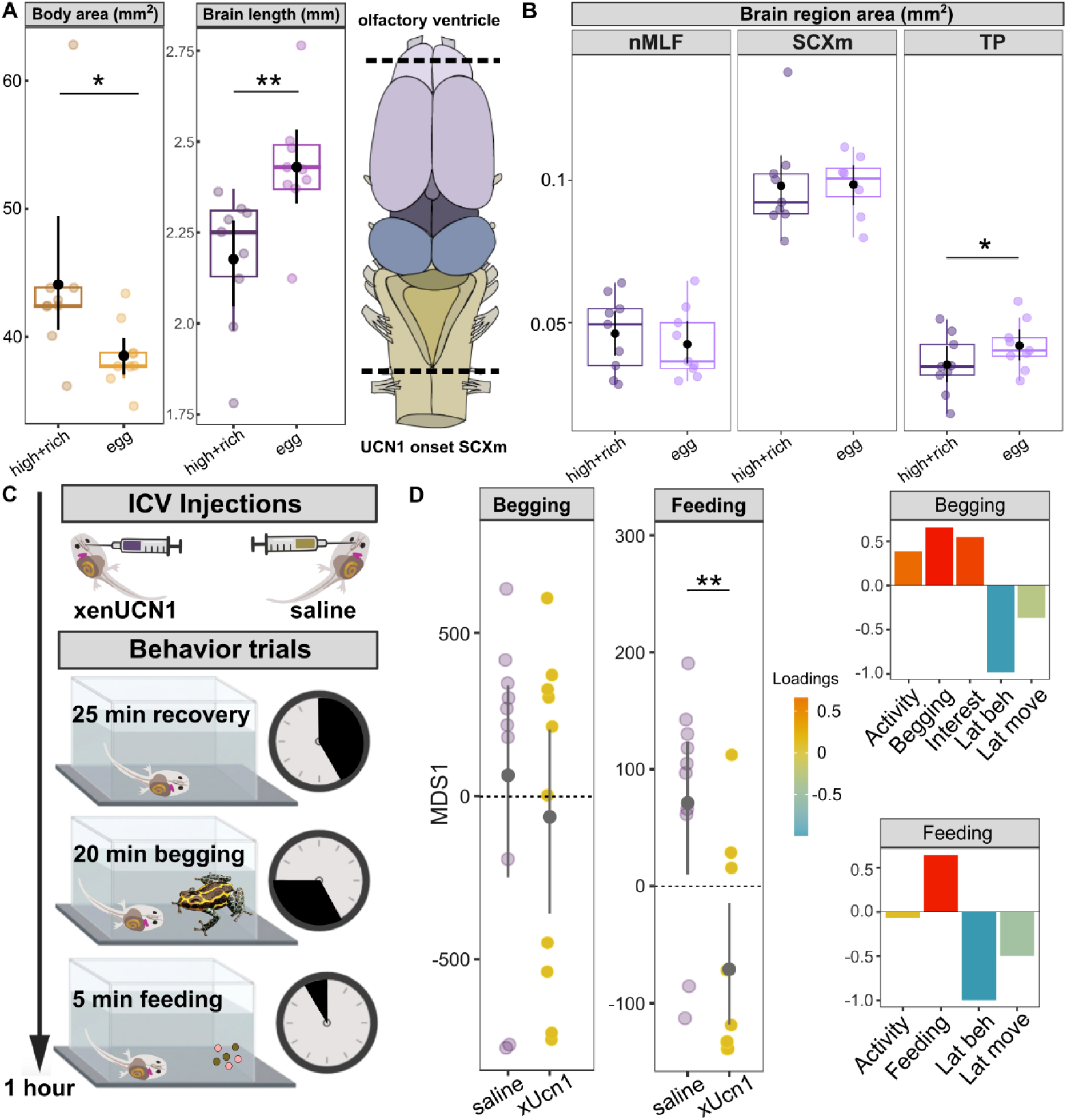
Dietary quality shapes growth of brain regions associated with begging behavior while Ucn1 suppresses feeding drive without promoting socio-positive behavior. **(A)** Body area and brain length along the anterior–posterior axis at stage 37 in tadpoles fed an enriched artificial diet or egg. Brain length was estimated using serial coronal sections between two neuroanatomical landmarks. Differences were tested using linear models. Brain schematic adjusted from (45). **(B)** Cross-sectional area of selected brain regions (nucleus of the medial longitudinal fasciculus (nMLF), spinal cord nucleus motorius nervi vagi (SCXm), posterior tuberculum (TP)) used to quantify region-specific brain growth. Differences were analyzed using a generalized linear mixed-effects model. **(C)** Experimental design for intracerebroventricular injection of Xenopus Ucn1 (xenUcn1) or saline followed by testing of begging and feeding behavior using each tadpole’s respective rearing diet. **(D)** Metric multidimensional scaling (mMDS) of behavioral responses during begging and feeding trials following saline or xenUcn1 injection. Points represent individuals; black dots indicate group means ± 95% confidence intervals. Insets show variable loadings on MDS1; treatment effects were tested using PERMANOVA, with significant differences indicated by asterisks.

Cross-sectional area of selected brain regions increased with age (χ²₁ = 4.26, p = 0.039) and differed among brain regions (χ²₂ = 983.57, p < 0.001), with a significant diet × brain region interaction (χ²₂ = 8.46, p = 0.015) (Fig. 4B). Egg-fed tadpoles showed selective enlargement of the posterior tuberculum (TP; p = 0.031), whereas the spinal cord nucleus motorius nervi vagi (SCXm; p = 0.277) and the nucleus of the medial longitudinal fascicle (nMLF; p = 0.597) were not significantly affected by diet. Notably, the posterior tuberculum contains the TH-expressing neuron population previously shown to be required for begging behavior in tadpoles (44).

### Ucn1 may facilitate social engagement by suppressing a competing drive

To determine the function of Ucn1 during begging, we injected amphibian Ucn1 or saline into the third ventricle adjacent to the nMLF and quantified tadpole responses toward an adult female frog and food in separate trials (Fig. 4C, D). The *R. imitator* urocortin transcript encoded a 149 aa prepropeptide as the longest ORF (Fig. S5), containing a single conserved CRF-family domain and exhibiting the canonical CRF α-helical fold. Xenopus Ucn1, the closest commercially available peptide, shares the same domain architecture without additional structural motifs and was therefore used for functional injections. We tested four alternative roles for Ucn1 signaling with distinct behavioral predictions: (1) if Ucn1 initiates socio-positive food solicitation, injections should increase begging behavior; (2) if it facilitates movement, general activity should increase without selectively enhancing begging; (3) if it gates begging by suppressing feeding independent of TH signaling, begging behavior should remain unchanged while feeding decreases; and (4) if it terminates food solicitation, begging behavior should decrease following Ucn1 injection. Behavior of tadpoles injected with amphibian Ucn1 did not differ from saline-injected controls in begging time or general activity levels (PERMANOVA: F₁,₁₈ = 0.69, *R*² = 0.04, *p* = 0.48) while feeding behavior was markedly reduced (F₁,₁₈ = 9.04, *R*² = 0.33, *p* = 0.004) (Fig. 4 D). Together with behavior-specific gene expression, these results indicate that Ucn1 reallocates internal motivational states by suppressing feeding drive, thereby permitting socio-positive food solicitation rather than initiating or terminating the behavior.

## Discussion

Early-life diet affects body growth, brain development, and behavior, yet most studies link diet to only one of these outcomes at a time and typically assess behavioral consequences in adulthood rather than in juveniles (28, 46). Here, we use poison frog tadpoles as a simple vertebrate model that allows us to link diet, neural development, and social behavior without constraints of intrauterine development or maternal dependency (30, 31, 47). By independently manipulating dietary quantity and quality and comparing artificial diets to a natural egg-based diet, we show that body growth scales primarily with food quantity, whereas developmental advancement more strongly associates with dietary quality. At the behavioral level, diet quality alone selectively shaped behaviors linked to parent–offspring interactions rather than social behavior broadly, with both begging and feeding behavior altered specifically in tadpoles reared on a natural diet. Together, these results highlight nutritional quality, rather than caloric intake alone, as a primary driver coordinating developmental progression and behavior in tadpoles.

Diet quality affected neural development at two mechanistic levels likely underlying the observed developmental and behavioral effects: first, through differential growth of specific brain regions rather than uniformly increasing brain size, expanding areas with TH^+^ populations mediating begging behavior; second, through selective modulation of specific neuron populations, altering the abundance of appetite-regulating Ucn1-and CART-expressing neurons but not TH-expressing populations. This suggests that dietary quality directly targets systems involved in internal state regulation, while affecting circuits in nutrition-sensitive regions through changes in regional brain volume rather than cell-specific modulation.

Diet can induce striking phenotypic plasticity in taxa (48–50), including anuran larvae (51–53). In our study, body and brain growth were dissociated, suggesting that diet regimes influence how resources are allocated along the larval developmental trajectory (54, 55). This phenomenon, termed “brain sparing”, is well described in the context of quantity, where restricted nutrition leads to preferential allocation of resources to the brain at the expense of body growth (56–58). Under such a model, reduced protein intake in egg-fed tadpoles would be expected to decrease body size while maintaining brain volume. However, this framework cannot account for the accelerated development and increased brain growth observed here. Instead, dietary quality likely drove these effects independently of macronutrient amount, potentially via bioactive compounds such as growth factors and hormones present in eggs but absent from artificial diets (59–61). Region-specific sensitivity to dietary variations was also reported in other systems (46, 55, 62, 63), with regions governing dopaminergic circuits selectively targeted in rodents (46). Notably, behavior was consistently affected in both systems, but differences were associated with risk avoidance and reaction to aversive stimuli in rodents and with differences in begging in anurans.

The selective expansion of anorexigenic neuronal populations under a natural diet is in line with evidence from mammalian studies of maternal obesity, where nutritionally imbalanced diets bias embryonic hypothalamic feeding circuits toward orexigenic drive (46, 59, 64). While most work has focused on canonical feeding peptides such as leptin and neuropeptide Y (65–71), our data highlight Ucn1 and CART as potent anorexigenic systems shaped by early diet. Ucn1 is a highly conserved member of the corticotropin-releasing factor (CRF) peptide family and is characterized by exceptionally high affinity for CRF receptors and binding proteins, exceeding that of CRF and other urocortins (72, 73). In the brain, Ucn1 shows locally restricted expression and overrides orexigenic signals such as orexin, positioning it as a candidate modulator of motivational state rather than solely a mediator of stress responses (74, 75).

In vertebrates, Ucn1 has been implicated in a range of behaviors beyond feeding, including reward processing, anxiety-related responses, maternal aggression and stress adaptation (73, 74, 76). Our findings extend the functional repertoire of Ucn1 by linking it to a socio-positive behavior in an early diverging vertebrate lineage, suggesting evolutionary co-option of this neuromodulatory system for distinct behavioral functions across taxa (9). Notably, although AgRP and orexin were upregulated during food solicitation, feeding behavior was reduced, consistent with a role for UCN1-associated pathways in counteracting these orexigenic signals during begging.

Together, diet-dependent modulation of Ucn1 neuron abundance, behavior-specific regulation of Ucn1 peptide availability, and functional tests under food-deprived conditions are consistent with a model in which Ucn1 does not initiate begging behavior but suppresses the competing drive to forage, possibly biasing motivational prioritization toward social engagement (77).

In an evolutionary context, prioritization of development over somatic growth may reflect adaptation to a parental care strategy that is extremely costly for parents, involving repeated investment of unfertilized eggs into offspring development rather than offspring production (36, 37). Efficient tuning of developmental, neural, and behavioral mechanisms that facilitate food solicitation in offspring may allow rapid progression to metamorphosis as shown in other systems (54, 78, 79), thus limiting the cumulative costs of prolonged provisioning for parents. How diet-induced developmental differences shape feeding drive and social behaviors after metamorphosis remains to be determined.

Our results suggest that nutritional quality can drive region-specific brain growth in anurans independent of food quantity, with direct consequences for early-life behavior. We further show that an evolutionary ancient anorexigenic neuromodulator is developmentally tuned by diet and recruited to suppress foraging drive during social engagement.

## Methods

### Model organism

*Ranitomeya imitator* is a small, diurnal poison frog (Family: Dendrobatidae) that forms stable pair bonds and exhibits obligate biparental care (34, 37). After hatching, males transport tadpoles to individual, small plant-held water bodies, which are characterized by extremely limited nutritional resources (35). Parents return repeatedly throughout larval development, with females provisioning tadpoles by depositing unfertilized trophic eggs that serve as their primary food source (36). Tadpoles are cannibalistic and exhibit a range of stimulus-specific behaviors. They attack conspecific tadpoles (aggression), approach and vigorously vibrate against adult frogs to solicit food (begging) and suppress movement and feeding in response to predators (freezing/ risk avoidance).

### Captive-bred animals

All tadpoles used in this study were produced by captive bred *Ranitomeya imitator* individuals within our laboratory poison frog colony. Adult frogs were maintained as breeding pairs (one adult male and one adult female) in glass terraria (45.72 × 30.48 × 30.48 cm; Exo Terra, Rolf C. Hagen USA, Mansfield, MA). Enclosures contained sphagnum moss substrate, driftwood, live *Pothos* plants, and horizontally positioned film canisters to provide egg deposition sites, as well as water-filled film canisters for tadpole deposition. Water was treated using a reverse osmosis conditioner (R/O Rx, Josh’s Frogs, Owosso, MI). Terraria were automatically misted ten times per day for 20 s per event. Frogs were fed live *Drosophila melanogaster* supplemented with vitamin powder and springtails three times per week. Housing was maintained on a 12:12 h light–dark cycle (lights on from 06:00 to 18:00). Temperature and relative humidity were recorded daily and typically averaged ∼25 °C and ∼95% within enclosures. Water-filled film canisters were inspected each morning for the presence of tadpoles. Upon detection, transported tadpoles were transferred to individual plastic cups containing 40 mL of water for experimental rearing and assigned a dietary regime. Siblings were assigned to different treatments to minimize confounding effects of shared genetic background. Water in all rearing cups was replaced three times per week, four hours after each feeding event.

### Feeding regimes

Tadpoles were reared in individual cups and assigned to one of five dietary treatments differing in food quality and quantity. Artificial diets consisted of a low-quality diet of ground dendrobatid tadpole pellets (Josh’s Frogs) (diet regimes B and D), commonly used in captive breeding, and a high-quality diet composed of ground protein-rich brine shrimp mixed with dendrobatid tadpole food at a 2:1 ratio (diet regimes A and C). In addition, tadpoles were fed poison frog eggs as a natural diet (diet regime E), representing the highest-quality food source, as trophic eggs provided by parents constitute an important food of *R. imitator* tadpoles in the wild. To enable standardized feeding with eggs of a similar developmental stage, fresh eggs from *Allobates femoralis* were used, as this dendrobatid species produces larger clutches (∼25 eggs) than *R. imitator* (1–2 eggs). Tadpoles receiving artificial diets were fed three times per week. Tadpoles in high-quantity treatments received 10 mg of food per feeding (30 mg per week, diet regimes A and B), an amount considered ad libitum as it was never fully consumed. Tadpoles assigned to low-quantity treatments received 2.5 mg per feeding (7.5 mg per week, diet regimes C and D), which was consistently consumed. Tadpoles in the egg-feeding treatment received two fresh *A. femoralis* eggs per week, corresponding to approximate feeding rates in the wild (1-2 eggs every 7.5 days, (37)). Based on an average dry mass of 2.475 mg per egg (after 6 h drying), the egg-based diet provided 4.95 mg dry food per week and was therefore classified as a low-quantity diet.

### Monitoring of tadpole ontogeny and behavioral trials

Tadpoles were reared to prometamorphic stage 37 (42) on their assigned diets, and body length, body width, and tail length were measured weekly with a caliper alongside developmental staging (Fig. S6). Animal sex cannot be determined at the tadpole stage. A subset of tadpoles displayed abnormal phenotypes under specific diets, including disproportionate lateral growth under the high-quantity base diet and arrested development (>270 days) under the low-quantity enriched diet (Fig. S6). As these phenotypes likely reflect a genuine effect of diet rather than a measurement artifact, affected individuals were retained in all analyses. Once individuals reached stage 36, they were marked and food-deprived for two days to standardize hunger. On the third day, each tadpole underwent a series of behavioral trials. After a 10-minute acclimation period in the testing arena, tadpoles were exposed to a bolt (non-social handling control), an adult female frog (to assess socio-positive food-soliciting behavior), and a smaller conspecific tadpole (to assess aggression) in randomized order. Behavior towards an inanimate object (a bolt) was assessed for 5 minutes to exclude individuals that responded affiliatively or aggressively toward a non-social stimulus. Each trial lasted 15 minutes, with tadpoles transferred to a clean arena and allowed to rest for 10 minutes between exposures. Feeding trials were conducted last to maintain a consistent energy state across all behavior assays. Feeding was recorded for 5 minutes, after which a well-fed *Dolomedes okefenokensis* (fishing spider) contained in a clear box was placed over the arena to measure predation threat-induced behavioral (feeding) inhibition for the next 5 minutes (Supplemental methods, Fig. S2). All trials were video-recorded and scored blind in BORIS (80) using a standardized ethogram (Table S3). After completing all behavioral assays, tadpoles were returned to their housing containers and fed according to their treatment group. Following an additional two-day fasting period, the socio-positive behavior assay was repeated to assess behavioral repeatability. Tadpoles were then anesthetized and sacrificed for tissue collection by submerging them in in frog water containing an MS-222 overdose.

### RNA–protein double staining

RNA–protein co-staining was used to validate Ucn1 antibody specificity (Ucn1 protein + Ucn1 mRNA) (Fig.S4B) and to confirm Ucn1 expression in neurons active during socio-positive, food-soliciting behavior (pS6 + Ucn1 mRNA). Combined immunohistochemistry and HCR in situ hybridization was performed on 15 µm brain cryosections using a protocol adapted from Molecular Instruments and the SHInE method (81, 82). A set of 20 HCR probes spanning the *Ranitomeya imitator* Ucn1 transcript (Molecular Instruments) was used for mRNA detection. Tissue sections were fixed in ice-cold 4% paraformaldehyde, dehydrated through graded ethanol, and processed without proteinase K treatment to preserve protein epitopes. Hybridization was performed using 1.6 µL of 1 µM probe set in 100 µL hybridization buffer, followed by fluorescent hairpin amplification (6 pmol each of hairpins h1 and h2 in 100 µL amplification buffer). Immunodetection of phosphorylated S6 (pS6; rabbit polyclonal, Catalog #44-923G) or Ucn1 (rabbit recombinant monoclonal, Catalog #ab283503) was carried out during the HCR amplification step using primary antibodies (1:200) and Alexa Fluor–conjugated secondary antibodies (1:500). Slides were mounted in DAPI-containing antifade medium and imaged the following day on a Leica DM6B fluorescence microscope at 20× magnification using Leica Application Suite X (LasX, v3.7.6.25997; Leica Microsystems).

### Immunohistochemistry

For immunohistochemical detection of Cocaine Amphetamine Related Transcript (CART) and tyrosine hydroxylase (TH) together with Ucn1, tadpole heads were fixed overnight at 4 °C in freshly prepared 4% paraformaldehyde (PFA) in 1× phosphate-buffered saline (PBS), washed three times in PBS, and cryoprotected overnight at 4 °C in 30% sucrose. Tissue was embedded in Tissue-Tek® O.C.T. compound and stored at −80 °C until cryosectioning. Brains were sectioned at 15 µm on a cryostat into two consecutive series and thaw-mounted onto SuperFrost Plus microscope slides, which were stored at −80 °C until staining. Fluorescent immunohistochemistry was performed as previously described (83). Sections were incubated overnight at 4 °C with primary antibodies diluted in blocking solution containing 5% normal goat serum, 0.3% Triton X-100 in 1× Tris-bufferd saline (TBS) (primary antibodies: Ucn1 [EPR25060-64] rabbit recombinant monoclonal antibody, Catalog # ab283503, Abcam, 1:500; CART 55-102 rabbit polyclonal antibody, Catalog # H-003-60, Phoenix Pharmaceutcals, Inc., 1:1500; TH: LC1 clone mouse monoclonal antibody Catalog # MAB318, Millipore, 1:500; PhosphoS6 (Ser244, Ser247) rabbit polyclonal antibody Catalog # 44-923G, Invitrogen, 1:2000). Following 3 washes in TBS, sections were incubated for 2 h at room temperature with species-appropriate Alexa Fluor–conjugated secondary antibodies (1:600) diluted in 10% normal goat serum in TBS (secondary antibodies goat anti-rabbit Alexa Fluor 594, goat anti-mouse Alexa Fluor 488, goat anti-rabbit Alexa Fluor 488, all highly cross adsorbed, ThermoFischer). Sections were rinsed in TBS and distilled water, coverslipped with Fluoroshield aqueous mounting medium containing DAPI (Catalog # ab104139, Abcam), and stored at 4 °C. Images were acquired on a Leica DM6B fluorescence microscope using a 20× objective and Leica Application Suite X software.

### Imaging and quantification of cell numbers and brain area

Images were acquired on a Leica DM6B fluorescence microscope using a 20× objective and Leica Application Suite X software. ImageJ (NIH, Bethesda, MD, USA) was used to count labeled neurons in both hemispheres and measure brain region area. We quantified cell number in the posterior tuberculum (TP), suprachiasmatic nucleus (SC), spinal cord nucleus motorius nervi vagi (SCXm), dorsal pallium (DP), and nucleus of the medial longitudinal fascicle (nMLF). Brain regions were delineated using a poison frog brain atlas developed for dendrobatids (84) and confirmed based on comparative descriptions (85, 86).

### Statistical analyses

All statistical analyses were conducted in R (v4.3.2; (87)). Data visualization was performed using the package ggplot2 (v4.0.0; (88)). All figures were compiled in Inkscape (v1.1; (89)).

Before model selection, we verified assumptions of normality and homogeneity of variance for all datasets using the Shapiro–Wilk test and either Levene’s test (package car, v3.1-2; (90)) or Bartlett’s test, as appropriate. Parametric analyses were applied when assumptions were met; otherwise, equivalent non-parametric alternatives were used. Where mixed-effects models were fitted, random structures were defined according to experimental design, and model residuals and dispersion were examined using DHARMa (v0.4.7; (91)) to confirm adequate model performance. Model selection, test statistics, and effect sizes are reported for each analysis in the results.

### Effects of diet on somatic growth

Tadpole body area at a stage close to metamorphosis (Gosner stage 36) was estimated as the area of an ellipse calculated from body length and width and compared among diet treatments to assess the effects of diet, food quantity, and quality. Differences between diet groups were first evaluated with a Kruskal-Wallis test followed by Benjamini-Hochberg-corrected pairwise Wilcoxon tests. To separate the effects of diet quantity and quality, tadpole area was log-transformed and analyzed using a Gaussian generalized linear mixed model (*glmmTMB*, v 1.1.8; (92)) including parental identity as a random effect.

### Effects of diet on developmental rate

Developmental time was quantified as the number of days required to reach Gosner stage 36. Differences among diet treatments were analyzed with Cox proportional hazards models with diet as a predictor using the package *survival* (v 3.5-7; (93)). Parental identity was included in the model as a stratification factor to account for genetic relatedness. The proportional hazards assumption was verified with the function *cox.zph*, and model fit was evaluated using likelihood ratio, Wald, and log-rank tests. To further assess the effects of diet quantity and quality, both predictors were included in a stratified model, and hazard ratios (HR) were estimated for each level relative to the reference category.

### Effectiveness of the newly established risk avoidance assay

To evaluate the effectiveness of the behavioral inhibition assay, we compared tadpole responses between trials with food alone and trials with food presented alongside a predator. Behavioral latencies were compared between conditions using paired Wilcoxon signed-rank tests for movement and feeding. Resulting *p*-values were adjusted for multiple testing using the Bonferroni correction.

### Behavioral data analysis

Behavioral data were analyzed separately for begging, aggression, feeding, and impulse-control trials. For each behavioral context, matrices were constructed with individual tadpoles as rows and behavioral parameters as columns (begging: activity, begging, interest, latency to behave, latency to move; aggression: activity, aggression, interest, latency to behave, latency to move; feeding: activity, feeding interactions, latency to feed, latency to move; risk avoidance: activity, feeding, latency to feed, latency to move). Differences in overall behavioral profiles among treatments were tested using permutational multivariate analysis of variance (PERMANOVA) based on Euclidean distances and visualized using metric multidimensional scaling (mMDS). Variable loadings on the first mMDS axis were used to determine the contribution of individual behavioral parameters to group separation.

### Repeatability of socio-positive behavior

To assess the reproducibility of behavioral responses, we retested socio-positive behavior in all tadpoles on a second day. Behavioral variables recorded during both sessions were reduced to one begging score using metric multidimensional scaling (mMDS) based on Euclidean distances. The first dimension (MDS1), explaining 74.4% of total variance, was positively associated with activity, affiliation, and interest, and negatively with latency to approach. Repeatability of the affiliation score across days was estimated using the rpt function from the rptR package (94), which calculates repeatability (R) from mixed-effects models by partitioning variance into between-and within-individual components. We fitted a model with individual ID as a random effect and obtained confidence intervals using 1000 bootstraps. To control for potential effects of nutritional history, we ran an additional model including rearing diet as a fixed effect.

### Phospho TRAP sample and data processing

We used phosphoTRAP to molecularly profile active neurons in tadpole brains under begging, aggressive, and control conditions. The control treatment consisted of a bolt to mimic stimulus addition without social interaction. Samples were processed and sequenced as described previously (43, 44, 84). In short, RNA was extracted from whole-brain tissue of five individual tadpoles per condition, and both total RNA (input) and the RNA fraction obtained by immunoprecipitation of phosphorylated ribosomal subunit S6 (pS6; IP) were sequenced for each sample. Raw paired-end reads from phosphoTRAP input and immunoprecipitated (IP) libraries were quality-filtered and trimmed using Trim Galore! (v0.5.0; (95)) with parameters optimized for Illumina data (--paired --phred33 --length 36 -q 5 --stringency 5 --illumina -e 0.1). Trimmed reads were pseudo-aligned to the *Ranitomeya imitator* transcriptome using Kallisto (v0.46.1; (96)) implemented within the Trinity framework (v2.8.4; (97)) with strand-specific library orientation. Gene-level abundance estimates were derived from transcript-level quantifications using Trinity and combined across all input and IP samples to generate a count matrix for downstream differential enrichment analyses. Gene-level count data from input and immunoprecipitated (IP) libraries were analyzed with a paired design to account for within-individual comparisons using the package edgeR (v3.42.4; (98)). Library sizes were normalized using the trimmed mean of M values (TMM) method. A generalized linear model (GLM) with a negative binomial distribution was fitted for each gene, including individual identity as a blocking factor and treatment condition as a fixed effect. Dispersion estimates were obtained with robust estimation, and likelihood ratio tests were applied to identify differentially expressed genes between conditions. Because behavior-related gene expression was assessed in genetically heterogeneous, non-clonal individuals, we applied a relaxed Benjamini–Hochberg false discovery rate threshold (adjusted p < 0.1) to retain sensitivity for biologically meaningful transcriptional changes. Genes with an absolute log₂ fold change greater than 1 were considered significantly differentially expressed. Annotated gene tables were cross-referenced against a curated list of orexigenic and anorexigenic neuropeptide systems to identify genes associated with energy sensing. Behavior-specific expression changes were distinguished from transcriptional responses related to handling or introduction of a novel object by excluding genes that were significantly regulated in the same direction in the control condition. Because only begging behavior was significantly influenced by diet, we restricted our search to neuronal populations specifically engaged in this behavioral context and excluded neurons that showed similar regulation in behaviors not affected by diet, such as aggression. In addition to the nutrition-focused analysis, we performed a complementary analysis to identify neuropeptidergic genes involved in behavioral regulation independent of their role in energy sensing. Significantly altered genes were cross-referenced against the Neuroactive ligand–receptor interaction KEGG pathway (hsa04080), augmented with neuropeptides from multiple taxa curated in the NeuroPep database (99). As for the primary analysis, genes showing significant regulation in the same direction in the handling/novel object control condition were excluded. Results of this broader neuropeptidergic analysis are provided in the Supplementary Material (Table S1, S2).

### Effects of artificial and natural high quality diet on brain development

To assess potential diet-related changes in brain morphology, we analyzed the brain and body of tadpoles from diets A and E, as these groups showed the most pronounced differences in begging behavior. Brain length and the area of major regions were quantified from serial 15 µm sections. Brain length was measured as number of cut sections between two neuroanatomical landmarks, the bulbus olfactorius (OB) and the beginning of the SCXm, marked by the occurrence of Ucn1 neurons in this area. All brains regions central to our hypothesis were included in the analysis of regional brain size (TP, nMLF, and SCXm). For each tadpole, mean brain area was calculated across slices for each brain region. Differences in regional brain areas were analyzed using a linear mixed-effects model (lmer, lme4 v1.1-35; (100)) with nutrition, brain region, and their interaction as fixed effects, age as a covariate, and tadpole ID as a random effect. Model terms were evaluated using Type II Wald χ² tests (car v3.1-2; (90)), and pairwise diet contrasts within brain regions were estimated using marginal means (emmeans v1.10.0; (101)) with Tukey adjustment. Brain length and tadpole area were analyzed separately using linear and Gamma models respectively, including nutrition and scaled age as predictors and model terms were evaluated with Likelihood Ratio (LR χ²) test.

### Effects of diet on abundance of three selected neuronal populations in the fore-mid-and hindbrain

To assess potential diet-related changes in brain morphology, we analyzed the brains of tadpoles from diets A (*n* = 9) and E (*n* = 9), as these groups showed the most pronounced behavioral differences. To test whether diet affected Ucn1 neuron abundance, we quantified Ucn1-positive neurons in the nMLF and SCXm across tadpoles from diets A and E. One tadpole from diet E was excluded because the SCXm region did not meet tissue integrity requirements. Counts were normalized to regional brain area (µm²) using a log offset in a negative binomial generalized linear mixed model (*glmmTMB*, v 1.1.8; (92)). The model included nutrition, brain region, their interaction, and scaled age as fixed effects, and tadpole ID as a random effect (nMLF and SCXm: 275 sections sections). Model significance was assessed using Type II Wald χ² tests, followed by Tukey-adjusted pairwise comparisons (*emmeans*, v 1.10.5; Lenth, 2024). To test whether nutritional effects on neuronal abundance were specific to Ucn1 or reflected a broader influence on anorexigenic circuits, we quantified CART-and TH-expressing neurons. CART neurons were counted in the nucleus of the medial longitudinal fascicle (nMLF: 143 sections) and TH neurons in the posterior tuberculum (TP: 206 sections). For both datasets, neurons were quantified from serial sections, and mean neuron counts per section were calculated for each tadpole. Generalized linear mixed models with a negative binomial error distribution and log link were used to analyze neuron counts (glmmTMB). Each model included nutrition and scaled age as fixed effects, tadpole ID as a random effect, and brain area as an offset term to correct for variation in measured tissue area.

### Ucn1 injection-induced behavioral differences

Behavioral variables were analyzed separately for begging and feeding trials. Begging and feeding behaviors were analyzed as described in the behavioral data analysis above. Differences in overall behavioral profiles between treatments (Ucn1 vs. saline) were then tested using a permutational multivariate analysis of variance (PERMANOVA) based on Euclidean distances and visualized using a principal coordinate analysis (PCoA; *pcoa* function, ape package; (102)) with loadings to determine which behaviors contributed most to separation along the first two axes.

### Quantification of dietary lipid and protein-levels

To quantify the nutritional composition of experimental diets and estimate nutrient intake, total lipid and protein contents were measured per gram of diet and combined with feeding frequency to calculate the amount of protein and lipid received over a two-week period. Measurements were performed on four biological replicates of artificial diets and frog eggs. Because tadpoles were fed fresh eggs from the poison frog *Allobates femoralis*, we additionally analyzed two eggs from *Ranitomeya imitator* to verify that egg nutritional content was comparable across species. Protein content overlapped between egg sources (*A. femoralis*: 51.1 ± 8.8 µg mg⁻¹, range 38.3–58.1, n = 4 replicates with 2 eggs each; R. imitator: 53.9 ± 15.2 µg mg⁻¹, range 43.2–64.7, n = 2 replicates with 2 eggs each). Lipid content in *A. femoralis* eggs (42.7 ± 6.8 mg g⁻¹, range 37.8–47.5, n = 2 replicates with 2 eggs each) fell within the range of the value measured for R. imitator eggs (39.6 mg g⁻¹, n = 1 replicate with 2 eggs) and these samples were therefore included in the analysis. To estimate nutrient intake for all diet regimes, per-gram lipid and protein values were multiplied by the total dry mass of diet provided over the two-week period for each feeding regimen. As tadpoles in high-quantity treatments did not fully consume the provided food at any feeding, whereas tadpoles in low-quantity and egg treatments consumed all provided food, nutrient intake for high-quantity diets was conservatively adjusted by dividing calculated values by two.

### Protein quantification

Protein was extracted by mechanical homogenization. Diet samples were homogenized in 0.8 ml cold phosphate-buffered saline (PBS) using a bead mill at 4 °C (speed 4.5, 10 cycles of 30 s with 10 s pauses). Homogenates were centrifuged at 2000 rpm for 5 min at 4 °C, and the clarified supernatant was used for protein quantification. Total protein concentration was determined using a Bradford colorimetric assay with bovine serum albumin (BSA) as a standard. Samples and standards were measured in triplicate in a 96-well plate, and absorbance was read at 595 nm. Protein content was calculated from standard curves and expressed relative to sample mass.

### Lipid quantification

Total lipid content was determined by petroleum ether–based gravimetric extraction. Diet samples were first oven-dried at 60 °C for 5 h to constant mass and weighed to obtain dry mass. Dried samples were then transferred to tubes, mechanically disrupted, and fully submerged in petroleum ether. Lipids were extracted over 8 days at room temperature, with solvent replaced every third day. After extraction, petroleum ether was removed, and samples were air-dried overnight followed by additional drying at 60 °C for 2–2.5 h. Lipid content was calculated as the difference between initial dry mass and lipid-free dry mass and standardized to dry weight.

### Intracerebroventricular injections into the third ventricle

Intracerebroventricular (ICV) injections were performed following established procedures adapted from (103). In short, injection needles were prepared by pulling borosilicate glass capillaries (3.5 inch length, 0.02 inch inner diameter; Drummond Scientific) using a P-97 micropipette puller (Sutter Instrument Co.) and trimming the tip under a dissecting microscope with fine forceps. Tadpoles were anesthetized in 0.04% buffered MS-222 and injected into the third ventricle using a Nanoject II microinjector (Drummond Scientific). The *R. imitator* urocortin sequence was retrieved from the NCBI RefSeq genome assembly aRanImi1.pri (GCF_032444005.1; Vertebrate Genomes Project, 2023) and comprised 1240 bp, with the longest predicted open reading frame (ORF4, NCBI ORFfinder) encoding a 149 amino acid prepropeptide. InterProScan detected a single conserved corticotropin-releasing factor (CRF) domain and no additional functional domains. AlphaFold structural prediction revealed the characteristic α-helical fold typical of CRF-family neuropeptides. The commercially available Xenopus Ucn1 peptide exhibited the same single CRF-domain architecture without additional predicted structural motifs. The conserved domain organization and structural similarity between R. imitator Ucn1 and Xenopus Ucn1 thus support functional equivalence. Therefore, Xenopus Ucn1 (xUcn1) was used for peptide injection experiments. Tadpoles (N = 10 per treatment, all reared on high quantity enriched diet) received either a saline control or Xenopus urocortin-1 (xenUcn1, NovoPro Bioscience Inc. Cat# Cat_312516, TFA removed) at a dose of 20 ng g⁻¹ body weight. The dose was chosen based on prior work demonstrating complete suppression of feeding behavior in *Xenopus* tadpoles (73) ensuring sustained peptide effects across the full behavioral testing period (1 h) despite the short half-life of Ucn1 (∼53 ± 3 min; (104)). All tadpoles resumed swimming within 4 min following injection. Behavioral testing commenced 20 min post-injection, beginning with a five minute baseline recordings, followed by an begging assay starting 30 min post-injection (20 min duration). Feeding behavior was assessed subsequently (five min duration) following a water change and a ten-minute inter-trial period. For begging trials, a non-maternal adult female was used as the stimulus individual.

## Permits

All experimental procedures were reviewed by the Institutional Animal Care and Use Committee (IACUC) at Stanford University (protocol 33097) and conducted in accordance with relevant guidelines and regulations.

## Data availability

Data files and data analysis scripts used to generate the results are available on DataDryad (pending public access) and GitHub (pending public access).

## Acknowledgements

We thank Madison Lacey for assistance with animal husbandry.

## Author Contributions

Conceptualization: MTF, LAO

Data Curation: MTF, BCG, LAO, MM

Formal Analysis: MTF, CRL

Investigation: MTF, LAO

Visualization: MTF, CRL, LAO

Methodology: MTF, BCG

Writing: MTF

Review and Editing: LAO, CRL, BCG

Project Administration: MTF, LAO

Supervision: LAO

Resources: LAO

Funding Acquisition: MTF, LAO

