## Supplemental material for "Early diet programs feeding circuits and food solicitation behavior"

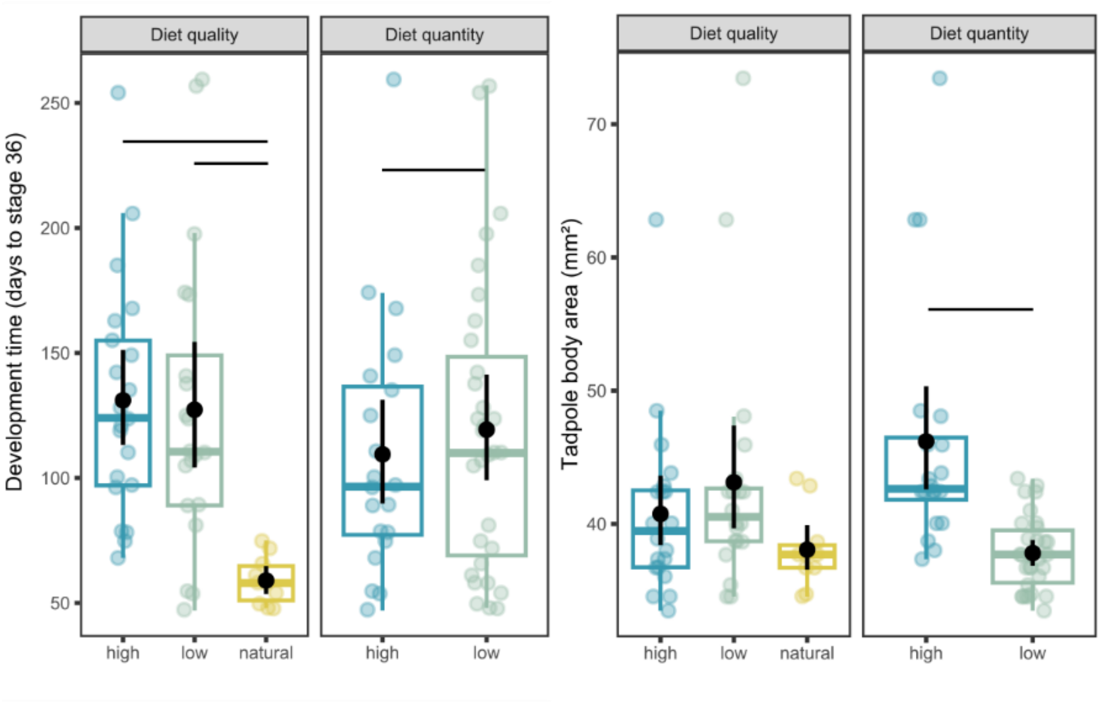


**Figure S1. Effects of diet quantity and quality on tadpole body size and developmental rate.** To assess diet quantity and quality effects, tadpoles were grouped by diet quantity (high: A, B; low: C, D, E) and diet quality (natural: E; enriched artificial: A, C; base artificial: B, D). **(A)** Tadpole body area at matched developmental stages (Gosner stage 36), analyzed using linear mixed-effects models with diet quantity and quality as fixed effects and parental identity as a random effect. **(B)** Developmental time to reach Gosner stage 36, analysed using Cox proportional hazards models including diet quantity and quality as predictors and parental identity used for stratification. Boxplots show median and interquartile range; points represent individual tadpoles and black dots indicate group means.

<u>Establishment of a risk avoidance assay</u>

To establish an assay that reliably measures predator-induced behavioral inhibition, we compared the latency to move and to feed in tadpoles exposed either to food alone or to food presented alongside a live spider. Each tadpole was tested sequentially under both conditions. Tadpoles showed markedly delayed responses when a predator was present: both movement latency ($V = 330$, $p = 0.002$, Bonferroni-corrected) and feeding latency ($V = 223$, $p < 0.001$, Bonferroni-corrected) were significantly longer in the spider condition compared to food alone. These results demonstrate that the assay effectively captures predator-induced freezing and thus represents a suitable framework for testing risk avoidance in tadpoles.

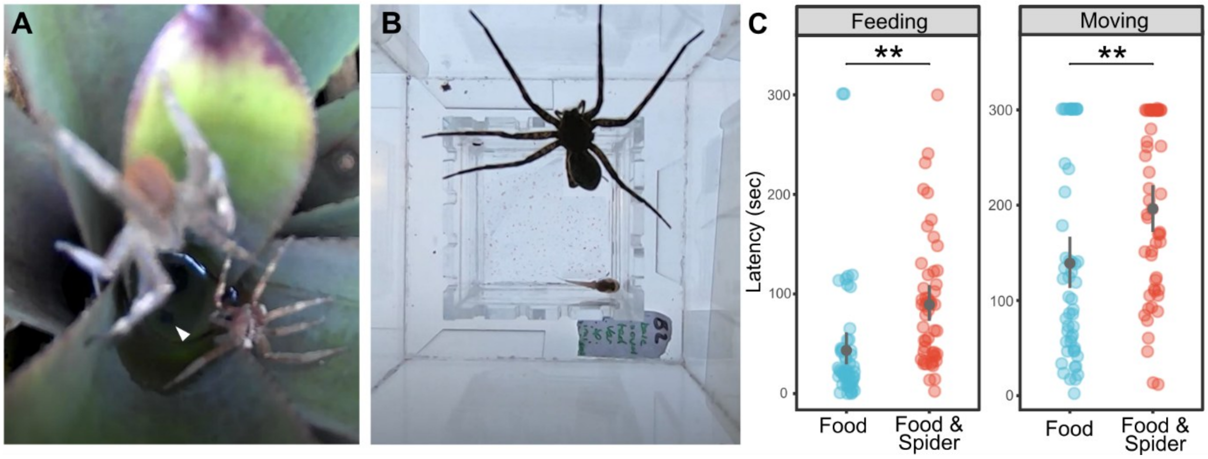


**Figure S2. Establishment of an assay for predator-induced behavioral inhibition. (A)** Natural predation context motivating assay design: a wandering spider of the genus *Phoneutria* approaches a tadpole nursery in plant leaf axils. **(B)** Experimental arena used to assay predator-induced behavioral inhibition. A second, larger arena was fitted over the tadpole arena. Tadpoles were exposed either to food alone without a predator present in the upper compartment, or to food presented adjacent a live, well-fed fishing spider (*Dolomedes okefenokensis*) placed in the upper compartment. This design allowed tadpoles to perceive the predator via multiple sensory cues (e.g. vibration and vision). Following a starvation period of 2 days each tadpole was tested sequentially under both conditions in a counterbalanced design. **(C)** Latency to initiate feeding (left) and movement (right). Latencies were compared using paired Wilcoxon signed-rank tests with Bonferroni correction. Points represent individual tadpoles (N = 50); black dots indicate group medians.

<u>Robustness of behavior measurements</u>

To assess the robustness of our behavioral measurements, we retested socio-positive behavior in all tadpoles three days apart and compared behavioral outcomes across trials. Metric multidimensional scaling (mMDS) of variables yielded two components that explained over 90% of the total variance. The first component (DIM1) explained 74.4% of the total variance and represented activity, begging, and interest positively and latency negatively, and was used as a begging score. Repeatability analysis based on mixed-effects modeling revealed significant repeatability of socio-positive behavior across testing days (R = 0.37, 95% CI = 0.09–0.62, $p < 0.01$). When controlling for rearing diet, some of the variance in the begging score was explained by diet, resulting in a lower but still significant repeatability estimate (R = 0.22, 95% CI = 0.03–0.46, $p < 0.05$). These findings indicate that begging behavior was reliably quantified for tadpoles across days.

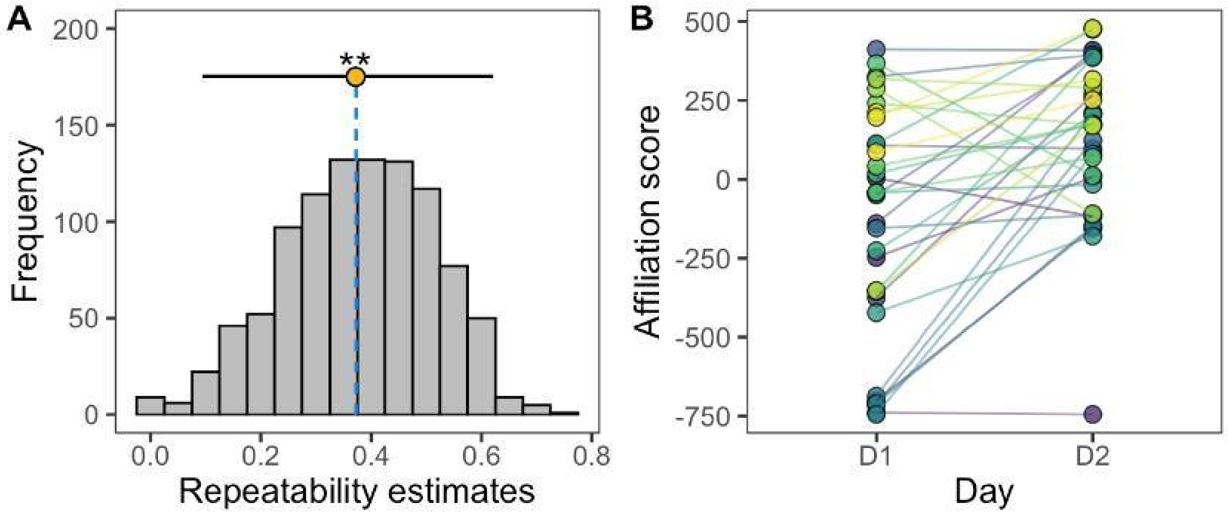


**Figure S3. Repeatability of begging behaviour across testing days.** Begging behaviour was quantified as described in Fig. 2 using a metric multidimensional scaling (mMDS)–derived begging score based on socio-positive behavioral variables. **(A)** Bootstrap distribution of repeatability estimates for tadpole identity based on mixed-effects models. The dot indicates the estimated repeatability and error bars denote 95% confidence intervals. **(B)** Individual begging scores measured across two testing days, with lines connecting repeated measurements of the same tadpole. Repeatability was estimated using the rpt function from the rptR package, with tadpole identity included as a random effect. Asterisks indicate significance ($p < 0.01$).

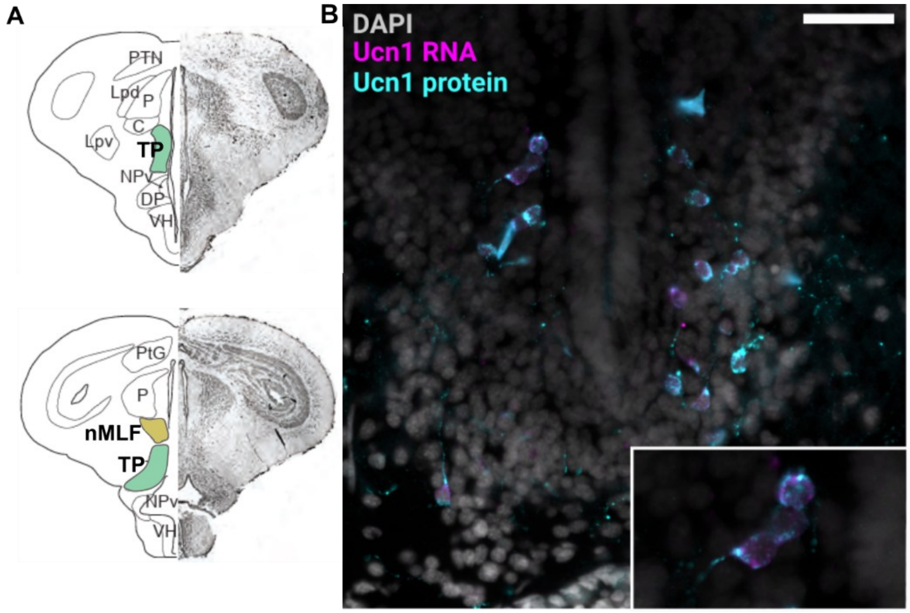


**Figure S4. Validation of Ucn1 antibody specificity by RNA–protein double staining.**
**(A)** Schematic brain sections adapted from (Fischer et al., 2019) indicating regions analyzed for quantification of TH+ (green) and Ucn1+ (yellow) neurons **(B)** Representative image of a 15 µm tadpole brain cryosection stained for Ucn1 mRNA (magenta; HCR *in situ* hybridization with a mix of 20 probes spanning the *R. imitator* Ucn1 gene, Molecular Instruments) and Ucn1 protein (cyan; IHC with rabbit recombinant monoclonal antibody, abcam), with nuclei counterstained with DAPI (gray). The inset shows a higher-magnification view of a single Ucn1-expressing neuron illustrating overlap between Ucn1 transcript and Ucn1 protein signal, indicating target specificity. Images were acquired on a Leica DM6B microscope using a 20× objective with a Leica K8 camera and Leica Application Suite X (LasX). Scale bar, 50 µm.

**A** *R. imitator UCN1 (NCBI RefSeq genome assembly aRanImi1.pri (GCF_032444005.1))*

>lcl|ORF4 (Longest predicted ORF, 149 aa)

MILTGQAGARPLERSFFEGPRLSLDGLLSDESAQALLYFIRERVLQMEVGDPQVQLRVSK
GSLSTEHEIPSQDLASMFSILRTVFQEFRPEESARTFLEVFSDRLKREDPPISIDLTYHILRH
MIDIAKTQNQKHQAEQNRIIFDSVGK

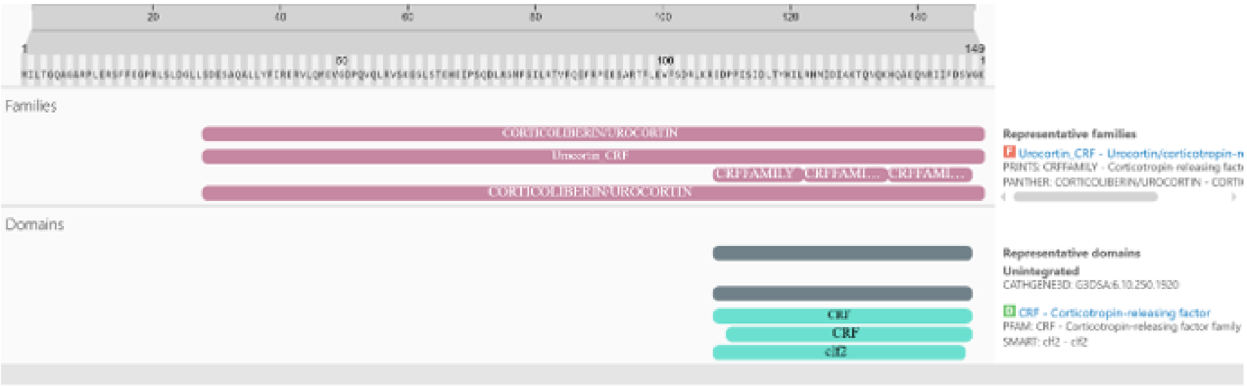


**B** *Xenopus UCN1* peptide (NovoPro, Cat# 3125)

EDPPISIDLTFHILRQMIEIAKTQNQKQQAEQNRIIFDSV

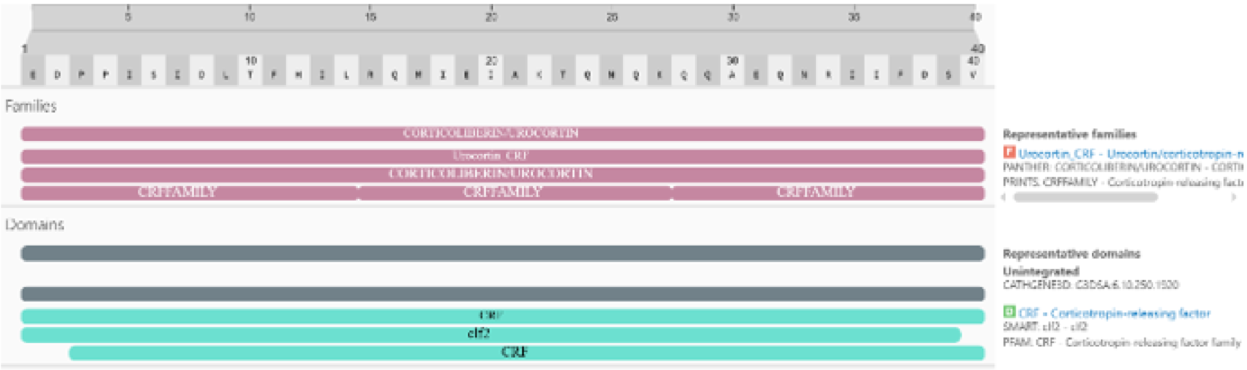


**C** Blast *Xenopus UCN1* vs *R. imitator UCN1*

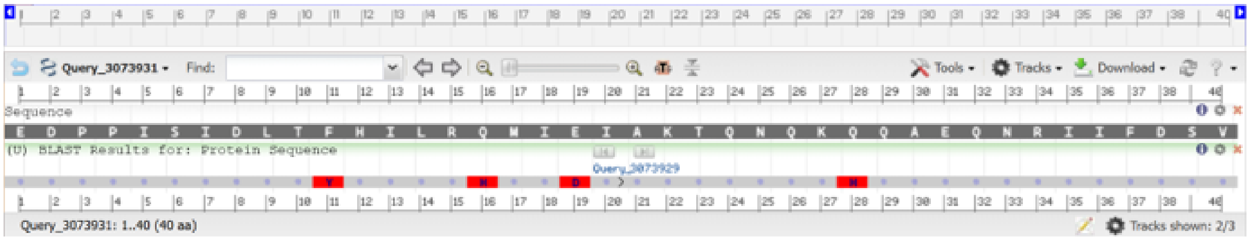


**Figure S5. Domain architecture and sequence comparison of R. imitator and Xenopus Ucn1 peptides.** **(A)** The *R. imitator* urocortin-1 sequence retrieved from the NCBI RefSeq genome assembly aRanImi1.pri (GCF_032444005.1). The longest predicted open reading frame (149 aa) encodes a prepropeptide. Domain analysis (InterProScan) identified a single conserved corticotropin-releasing factor (CRF) family domain and no additional functional domains. **(B)** Domain analysis of the commercially available *Xenopus* Ucn1 peptide used for injections (NovoPro, Cat# 3125). **(C)** Alignment of the *Xenopus* Ucn1 peptide (upper sequence) with the corresponding C-terminal region of the *R. imitator* prepropeptide (lower sequence)

**Table S1. Begging-specific neuropeptidergic gene regulation.**

Annotated gene lists were cross-referenced against curated orexigenic and anorexigenic neuropeptide systems and an expanded neuropeptidergic dataset derived from the Neuroactive ligand–receptor interaction KEGG pathway (hsa04080) and the NeuroPep database. Only genes showing begging-specific regulation were retained, defined as genes not significantly enriched and regulated in the same direction in either the handling/novel object control trial or the aggression trial. The table reports adjusted $p$-values ($p$adj; significance threshold < 0.1), enrichment status in the phospho-TRAP immunoprecipitated (IP) fraction, and $\log_2$ fold change ($\log_2$FC; significant > 1) relative to the total RNA fraction.

| Gene.Abbr | Gene.Name | category | logFC_begging | padj_begging | logFC_control | padj_control | enriched in control | logFC_aggression | padj_aggression | enriched during aggression |
|---|---|---|---|---|---|---|---|---|---|---|
| UCN1_MOUSE | Urocortin | Urocortin | 1.78 | 0.08 | -1.40 | 0.59 | no | -0.29 | 0.92 | no |
| OREX_CANLF | Orexin | Orexin | 1.57 | 0.04 | -0.77 | 0.75 | no | 1.44 | 0.27 | no |
| PDYN_PIG | Proenkephalin-B | PDYN | 1.48 | 0.01 | 2.21 | 0.13 | no | 1.33 | 0.27 | no |
| PTH_MOUSE | Probable peptidyl-tRNA hydrolase | PTH | 1.26 | 0.03 | 0.39 | 0.86 | no | 0.58 | 0.69 | no |
| CALCA_CHICK | Calcitonin gene-related peptide | Calcitonin gene-related peptide | 1.18 | 0.07 | 0.76 | 0.46 | no | 0.93 | 0.48 | no |
| NUCB2_HUMAN | Nucleobindin-2 | Nucleobindin-2 | 1.11 | 0.01 | 1.45 | 0.32 | no | 0.65 | 0.49 | no |
| CBLN1_MOUSE | Cerebellin-1 | Cerebellin-1 | 1.04 | 0.01 | 0.82 | 0.18 | no | 0.41 | 0.67 | no |
| P2RX5_HUMAN | P2X purinoceptor 5 | P2RX5 | -1.42 | 0.10 | -0.70 | 0.62 | no | -1.46 | 0.61 | no |
| PRL_CHICK | Prolactin | Prolactin | -1.86 | 0.04 | -1.12 | 0.26 | no | -2.26 | 0.17 | no |
| CRHBP_XENLA | Corticotropin-releasing factor-binding protein | Corticotropin | -2.99 | 0.05 | -2.13 | 0.18 | no | -0.30 | 0.94 | no |
| VIP2_MOUSE | Inositol hexakisphosphate and diphosphoinositol-pentakisphosphate kinase 2 | Inositol | -4.06 | 0.07 | NA | NA | NA | -2.63 | 0.68 | no |
| GLR_MOUSE | Glucagon receptor | Glucagon | -4.75 | 0.10 | -2.76 | 0.33 | no | NA | NA | NA |
| SV2C_RAT | Synaptic vesicle glycoprotein 2C | Glycoprotein | -5.15 | 0.07 | NA | NA | NA | -0.73 | 0.88 | no |
| IRS2A_XENLA | Insulin receptor substrate 2-A | Insulin | -5.64 | 0.01 | -2.48 | 0.33 | no | NA | NA | NA |
| GLRA3_RAT | Glycine receptor subunit alpha-3 | GLRA3 | -6.04 | 0.07 | NA | NA | NA | -2.77 | 0.69 | no |
| CD3G_BOVIN | T-cell surface glycoprotein CD3 gamma chain | Glycoprotein | -6.44 | 0.07 | NA | NA | NA | NA | NA | NA |
| IGF2B_XENLA | Insulin-like growth factor II-B | Insulin | -7.33 | 0.01 | -4.46 | 0.18 | no | NA | NA | NA |
| HRH3_CAVPO | Histamine H3 receptor | HRH3 | -7.40 | 0.06 | NA | NA | NA | NA | NA | NA |

**Table S2. Aggression-specific neuropeptidergic gene regulation.**

Genes associated with neuropeptidergic signaling were identified using the Neuroactive ligand–receptor interaction KEGG pathway (hsa04080) supplemented with neuropeptides curated from the NeuroPep database. Only aggression-specific genes were included, defined as genes not significantly enriched and regulated in the same direction in the handling/novel object control trial or the begging trial. Reported values include adjusted $p$-values ($p$adj; significance threshold < 0.1), enrichment status in the phospho-TRAP immunoprecipitated (IP) fraction, and $\log_2$ fold change ($\log_2$FC; significant > 1) relative to the total RNA fraction.

| Gene.Abbr | Gene.Name | category | logFC_tad | padj_tad | logFC_bolt | padj_bolt | enriched in control | logFC_female | padj_female | enriched during begging |
|---|---|---|---|---|---|---|---|---|---|---|
| SERPH_CHICK | Serpin H1 | Serpin | 1.72 | 0.09 | 0.50 | 0.48 | no | 0.41 | 0.36 | no |
| GALR2_HUMAN | Galanin receptor type 2 | Galanin | -7.40 | 0.00 | NA | NA | no | -5.13 | 0.15 | no |
| P2RX7_MOUSE | P2X purinoceptor 7 | P2RX7 | -7.95 | 0.01 | NA | NA | NA | NA | NA | NA |

Supplemental methods

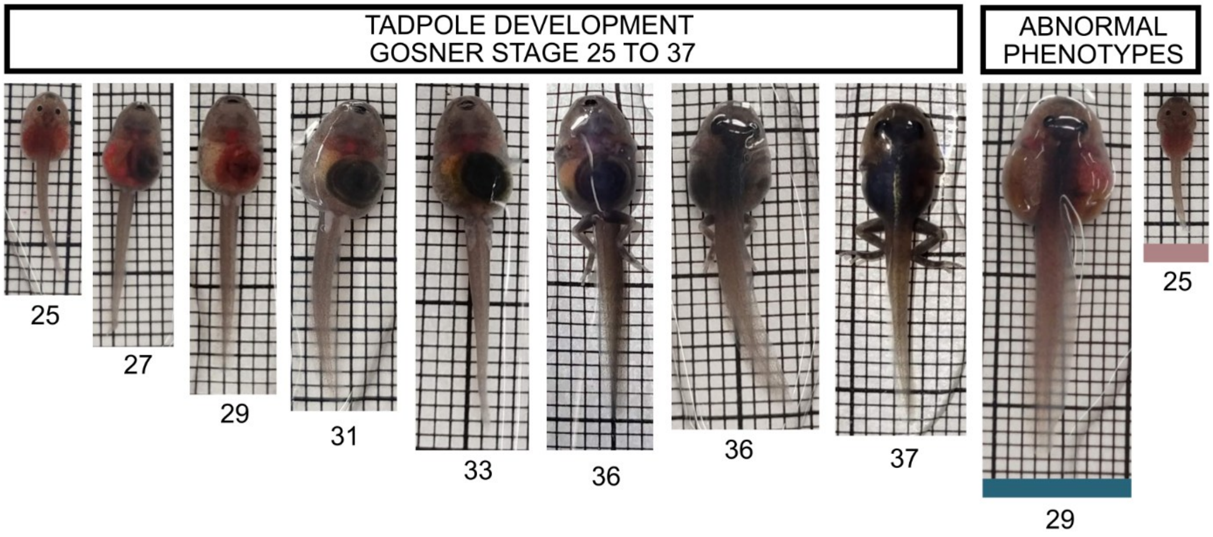


**Figure S6. Tadpole staging and abnormal phenotypes** Developmental staging (left panel) was based on toe differentiation and development following Gosner, where stage 25 corresponds to larvae after hatching without legs and all toes are separated at stage 37. Representative pictures of selected stages are shown. Abnormal phenotypes (right panel) were observed under the high-quantity base diet (blue; disproportionate lateral growth at an early stage) and the low-quantity enriched diet (pink; arrested development for more than 270 days).

Ethogram for behavior codings

Behavior was scored from video recordings using BORIS event-logging software (Friard & Gamba, 2016). Behaviors were coded as point (< 1 second) or state events (duration >1 second) and assigned to functional categories (activity, begging, aggression, fear, food, interest, or latency). Activity included discrete movement events, swimming bouts, and non–stimulus-directed wiggling. Begging behavior comprised stimulus-directed wiggling or begging within one body length of the stimulus and nipping events. Aggression included discrete aggressive bouts and sustained fights initiated by the focal tadpole, while fear-related behaviors captured instances in which the focal tadpole was attacked or aggressed upon. Feeding was scored as feeding states and feeding bouts, interest as time spent within one body length of the stimulus, and latency variables as the time from trial onset to the first movement or first affiliative, aggressive, or feeding behavior.

Table S3. **Ethogram used for coding of tadpole behavior.**

| **Behavior** | **Behavior type** | **Description** | **Category** |
|---|---|---|---|
| Move | Point event | Moving bout | Activity |
| Swimming | State event | Swimming | Activity |
| Begging | State event | Wiggling or begging towards stimulus (start or end within one body length of stimulus) | Affiliation |
| Nip | Point event | Nipping on stimulus | Affiliation |

| | | | |
|---|---|---|---|
| Aggressing bout | Point event | Events of bites, tackles, side swipes or displacement of conspecific | Aggression |
| Fight initiated | State event | Attacks conspecific resulting in longer lasting fight | Aggression |
| Is attacked | State event | Is attacked | Fear |
| Latency move | State event | Time until first move after starting trial | Latency move |
| Eating | State event | Feeding | Food |
| Proximity | State event | Closer than one bodylength to stimulus | Interest |
| Wiggling not mom | State event | Not stimulus directed wiggling | Activity |
| Latency behavior | State event | Time until first begging/aggression/feeding happens | Latency behavior |
| Is aggressed on | Point event | Is tackled, displaced or bitten by stimulus tad | Fear |
| Feeding bout | Point event | Short feeding event | Food |
